# Simulating CD8 T Cell Exhaustion: A Comprehensive Approach

**DOI:** 10.1101/2024.09.06.611697

**Authors:** Andrea J. Manrique-Rincón, Ben Foster, Stuart Horswell, David A. Goulding, David J. Adams, Anneliese O. Speak

## Abstract

Immunotherapy has revolutionised the treatment of multiple cancer types, however, these treatments only work for a proportion of patients and biomarkers to predict response are lacking. One correlate of response is the reinvigoration of a subset of CD8 T cells that have an exhausted phenotype and impaired functionality. In order to develop new therapies, reproducible models are required to identify candidate target genes that enables the reversal of key hallmarks of T cell exhaustion. Here we describe the development of an *in vitro* model by chronically stimulating T cells with their cognate antigen and performed an in depth temporal phenotypic characterisation. This model recapitulates many of the critical hallmarks of exhaustion, including increased expression of canonical exhaustion surface markers, impaired proliferation, reduced cytokine production, decreased release of cytotoxic granules, and metabolic alterations, including dysfunctional mitochondria. These exhaustion hallmarks were validated using an *in vivo* model and a gene signature identified which robustly define the shared *in vitro* and *in vivo* exhausted state. Critically, this signature is also observed in tumour infiltrating T cells from multiple human tumour types, validating the translational potential of this model for discovering and triaging new therapies.

## Introduction

Exhausted T cells emerge when their antigen remains present for long periods, impairing immune responses including viral infections and cancer ^1^. These cells were initially described after mice were infected with lymphocytic choriomeningitis virus (LCMV) clone 13 strain, which results in a chronic persistent infection ^2–4^. Exhausted T cells are characterised by an impairment in functionality, proliferation and metabolic alterations. This is accompanied by high expression of inhibitory receptors (IR) including programmed cell death protein 1 (PD1), cytotoxic T-lymphocyte association protein 4 (CTLA4), T cell immunoglobulin and mucin domain-containing protein 3 (TIM3), lymphocyte activation gene-3 (LAG-3), CD39, CD244 (2B4) and TIGIT ^5^. These changes can be identified at the transcript level and via epigenetic alterations^6^.

While it is evident that tumour antigen reactive T cells are present within tumours their ability to eliminate cancer cells is limited. This is because the tumour microenvironment (TME) provides several conditions besides chronic antigen exposure to maintain the exhausted phenotype of T cells, including; hypoxia ^7–9^, lack of nutrients ^10,11^, immunosuppressive cells like regulatory T cells ^12^ and Myeloid-derived suppressor cells (MDSC) ^6^, and the expression of inhibitory ligands such as PD-L1, and PD-L2 ^13–15^. These signals trigger and maintain tumour reactive T cells in an exhausted state exemplified by decreased proliferative capacity, reduced effector function, expression of inhibitory receptors (IRs) and metabolic changes including an increase in ROS and an increase of mitochondria mass with a decrease in oxygen consumption ^1,16–18^.

The success of therapies that boost the immune system to respond against cancer cells (immunotherapies) has brought a revolution characterized by a drive to identify new ways to manipulate different components of the immune response in order to avoid immune escape, and ultimately to eradicate tumours. However, an essential factor that hinders this potential is the exhausted phenotype of tumour-reactive cells ^19,20^. One of the most successful immunotherapies in clinical use is Immune Checkpoint Blockade (ICB). This therapy disrupts the interaction between IRs on exhausted cells and their ligands ^21–23^. However, only a fraction of patients will respond and many will suffer significant adverse events. Despite multiple clinical studies, biomarkers to predict patient response or risk of adverse events to ICB have not been fully characterised. Temporal investigations to identify the kinetics and identity of the cells that respond to ICB, have shown that only a subset of exhausted cells improved their functionality after ICB administration ^24–27^. Furthermore, ICB does not reverse the epigenetic changes that occur in the exhausted T cells and thus cells can revert after treatment cessation. In addition to ICB, another major type of immunotherapy is adoptive transfer of tumour reactive cells, in particular engineered chimeric antigen receptors T cells (CAR-T). CAR-T cells have shown remarkable responses in many haematological malignancies, yet their application to the treatment of solid tumours has been hindered by the suppressive tumour microenvironment which rapidly converts the cells into an exhausted state ^28–31^. With the advances in cellular manufacturing it could be possible to engineer exhaustion resistant CAR-T if suitable genetic targets were identified.

To better understand how to improve the reactive capacity of exhausted cells and harness it for immunotherapy, it is necessary to construct robust and reliable models of T cell exhaustion with translational potential. Here, we develop an *in vitro* model that uses exposure to their cognate antigen, mimicking chronic antigen exposure. This *in vitro* model reproduces the main characteristics of exhausted T cells and was validated using cells isolated from tumour-infiltrating lymphocytes (TILs). RNAseq of the *in vitro* and *in vivo* cells identifies shared gene signatures specifically associated with exhausted T cells. To confirm the translational potential of this model we compared the shared exhaustion signature we have identified from our models to a human pan-cancer T cell atlas ^32^ generating a list of 234 genes that are common between the models.

## Results

### A model of exhaustion *in vitro* that recapitulates the main hallmarks of exhaustion upon chronic stimulation

Exhausted T cells are produced via chronic stimulation through the T cell receptor (TCR) with cognate antigen. To recapitulate this process *in vitro*, we used cells from OT-1 mice carrying a transgenic TCR that recognizes the H-2K(b)/ SIINFEKL OVA (257-264) epitope ^33^. Freshly isolated CD8 T cells from spleens of OT-1 mice were stimulated every 48h in the presence of IL-2 and after 96h supplemented with IL-15 (Fig. 1A). After seven stimulations, these cells were analysed in comparison to early activated T cells (24 hours with SIINFEKL), and to freshly isolated naïve T cells. The *in vitro* exhausted cells showed a reduced ability to proliferate, compared to activated T cells, as evidenced by a reduction in cells in the S phase, determined by EdU incorporation (Fig. 1B). The expression of hallmark IRs including 2B4, CD39, TIM3, PD1, LAG-3 and CTLA4 was assessed. Expression of all these markers was higher in exhausted T cells (Fig. 1C,D). TIM3, CD39 and 2B4 were particularly highly expressed on the exhausted cells relative to activated cells enabling their use to distinguish these populations. Notably other frequently used exhaustion markers such as PD1, LAG-3 and CTLA-4 were also highly expressed by activated cells. Another critical hallmark of exhausted T cells is their reduced functional capacity regarding cytokine production and cytotoxicity. The *in vitro* exhausted cells contained higher levels of Granzyme B, a key component of cytotoxic granules, consistent with the phenotype reported in CD8 TILs ^24^ (Fig. 1E). The functional impairment of the *in vitro* exhausted cells was evidenced by the reduced polyfunctionality of the cells after restimulation with SIINFEKL, either the ability to degranulate and produce cytokines (CD107a+IFNγ+ or CD107a+ TNFα+) or to produce two cytokines simultaneously (IFNγ+ TNFα+) (Fig. 1F). Looking at the pool of exhausted cells and compared to the four-cell-stage developmental framework of exhaustion^34^, defined by Ly108 and CD69, these *in vitro* exhausted T cells lie between intermediately and terminally exhausted (Fig. S1), which allows the exploration of the development and diversity of exhausted cell populations *in vitro*.

**Figure 1.**
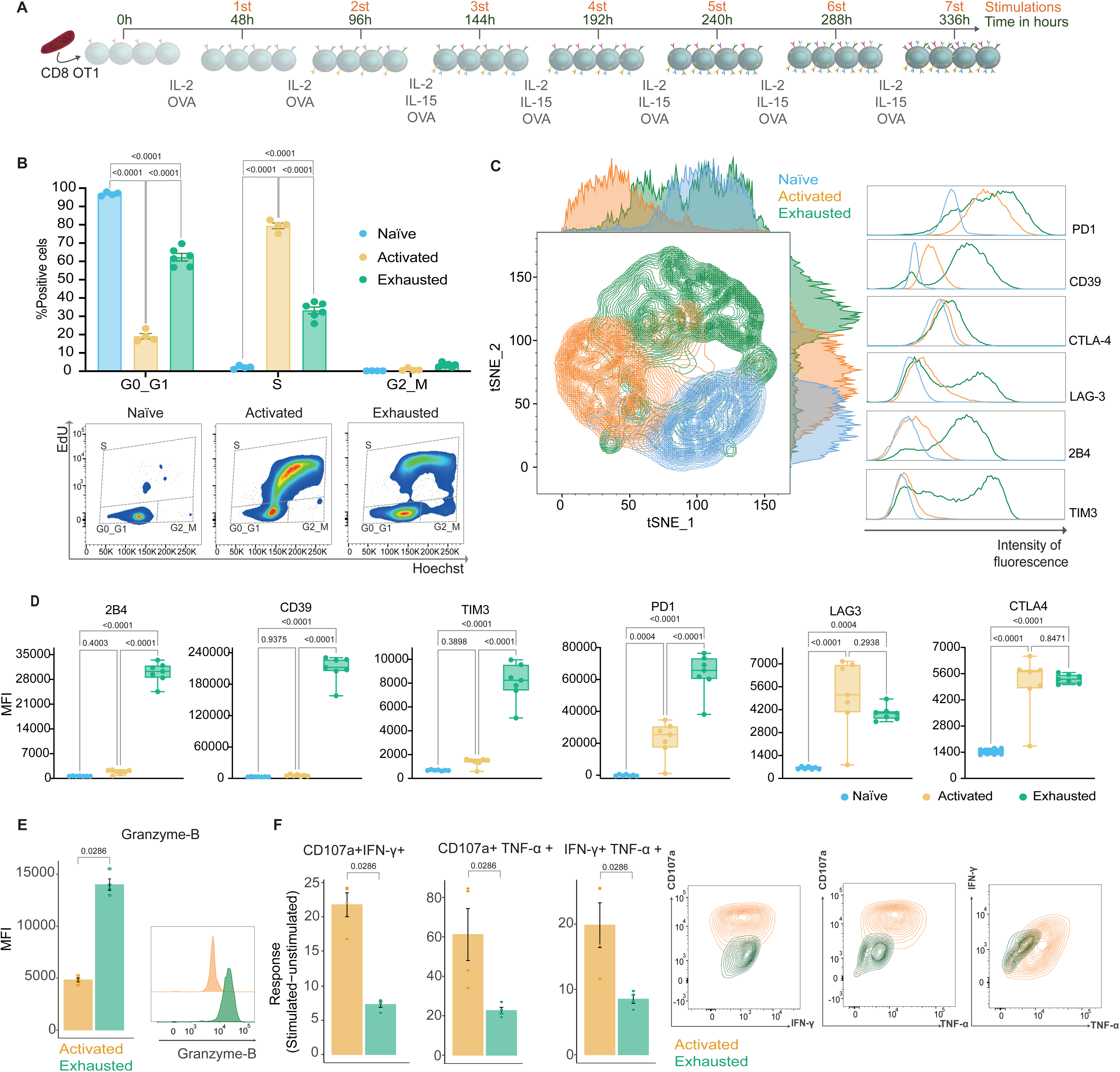
Characterisation of the development of *In vitro* exhausted CD8 T cells. Freshly isolated cells (Naïve), cells activated with SIINFEKL for 24h (Activated), and chronically stimulated cells (Exhausted) were phenotypically characterised. A. *In vitro* experimental design of chronic stimulation to produce exhausted T cells. B. Representative gating strategy and percentages of T cells per cell-cycle stage obtained by EdU and Hoescht staining. C. tSNE using IRs with representative histograms for indicated markers on naïve, activated, and *in vitro* exhausted T cells. D. Quantification of geometric mean fluorescence intensity (MFI) of individual markers. E. Intracellular expression of Granzyme B with representative histograms. F. Background unstimulated corrected percentage of double positive populations of CD107a+ IFNγ+, CD107a+ TNFα+, and IFNγ+ TNFα+ in stimulated cells with representative contour plot of stimulated samples. Representative data from three independent experiments. Mean ±SEM shown with symbols representing individual mice. Statistical analysis performed by two way ANOVA with a Tukey’s multiple-comparisons test (B,D) or Mann whitney tests (E,F) with significant p values indicated in the figure.

### *In vitro* exhausted cells have an altered metabolism and higher numbers of mitochondria

There is a change in the metabolism of T cells after the first encounter with an antigen. This event triggers a conversion from mitochondria-dependent oxidative phosphorylation (OXPHOS) to aerobic glycolysis, and once the effector stage has passed, memory cells return to a more quiescent state dictated by OXPHOS^17^. Chronic stimulation leads to disruption in the metabolism of exhausted compared to activated cells in multiple ways. The spare respiratory capacity (SRC) is a key factor in determining a cell’s response to stress and has previously been shown to vary between different CD8 subsets with a low SRC in effector cells and high SRC in memory. We determined the SRC of the *in vitro* exhausted T cells compared to activated and observed a significant reduction in the SRC of exhausted cells (Fig. 2A) accompanied by a reduced extracellular acidification rate (ECAR) (Fig. S2). Exhausted cells showed increased glucose uptake (Fig. 2B) with higher levels of oxygen species (ROS) (Fig. 2C). The exhausted cells had increased mitochondrial mass, as determined by mitotracker (Fig. 2D,E), and an increase in the mitochondrial membrane potential (Fig. 2F). In order to quantify mitochondrial number and morphology at an ultrastructural resolution we performed transmission electron microscopy (TEM) (Fig. 2G). Mitochondrial content of activated CD8 T cells is twice that observed in naïve CD8 T cells, with exhausted cells having a further two-fold increase compared to activated cells (Fig. 2G, H). However, the mitochondria of activated cells are bigger than those in exhausted and naïve cells (Fig. 2I) although their circularity was maintained (Fig. 2J). While both activated and exhausted cells are larger than naïve by 3.5 fold (Fig. 2K), the exhausted cells showed a reduction in their circularity and a more irregular shape (Fig. 2G,L).

**Figure 2.**
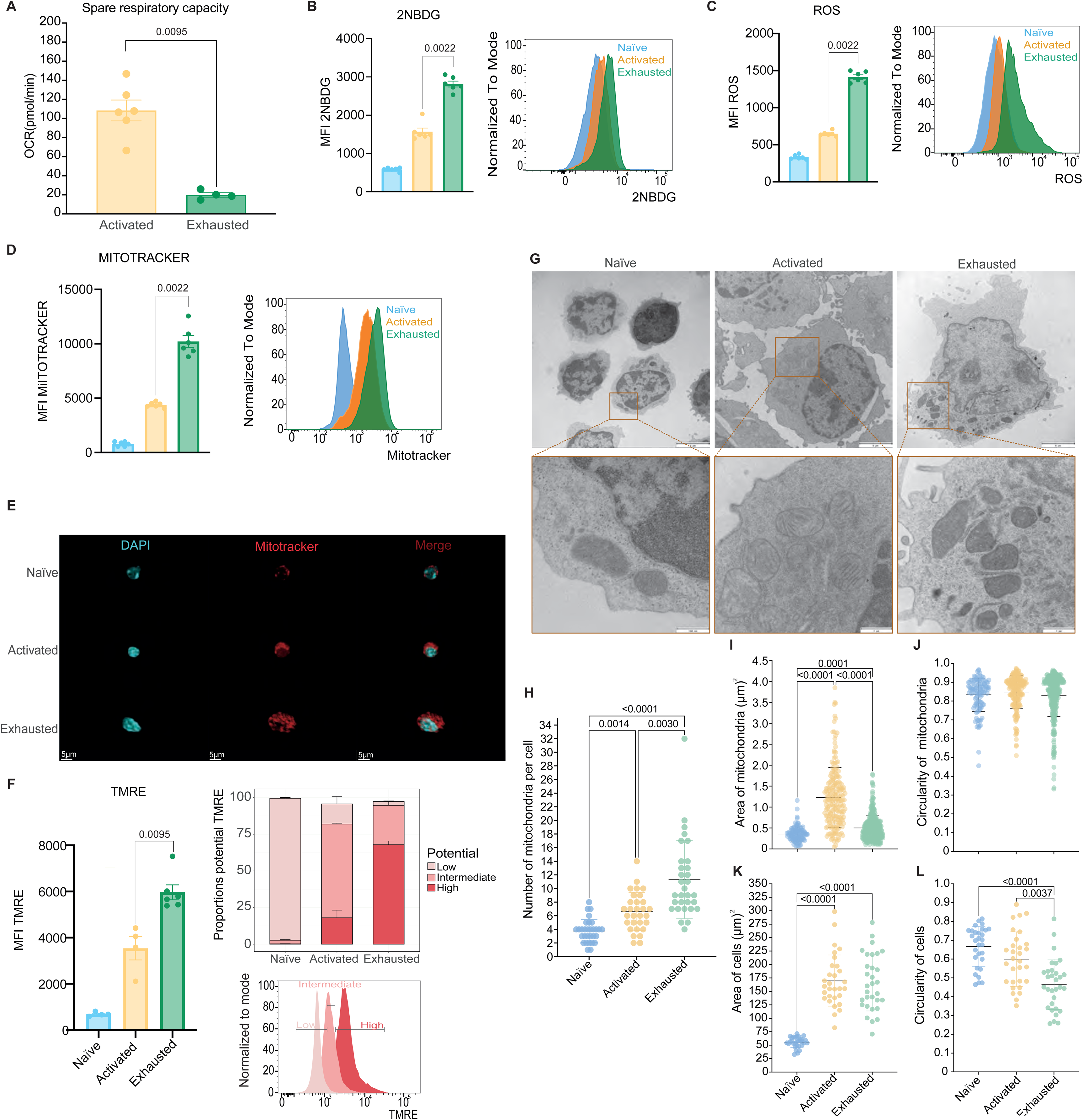
Metabolic alterations of *in vitro* exhausted T cells. Freshly isolated cells (Naïve), cells activated with SIINFEKL for 24h (Activated), and chronically stimulated cells (Exhausted) characterised for metabolic changes. A. Quantification of the spare respiratory capacity B. Glucose uptake quantification using 2-NBDG, representative histograms are shown. C. Determination of cellular ROS content with representative histograms shown. D. Measurement of mitochondrial mass with mitotracker deep red FM, with representative histograms shown. E. Confocal visualisation of mitochondria, 3D view with mitotracker in red and DAPI for nucleus in blue. Scale bar = 5μm. F. Mitochondrial membrane potential assessed using TMRE, representative histograms with gates of potential. G. Transmission electron microscopy (TEM) of naïve, activated, and exhausted T cells with inset zoom on mitochondria, scale bar as indicated in the panels. H. Number of mitochondria per cell. I-L. Area and circularity of mitochondria and cells respectively. Representative data from three (A-E) and two (F) independent experiments For TEM at least three sample preparations per group were imaged and at least 27 cells per sample were quantified. Mean and ±SEM shown with symbols representing individual mice (A-F), individual cells (H, K, L) or individual mitochondria (I, J). Statistical analysis performed by two-tailed Student’s t-tests (A-D,F) and Kruskal-Wallis ANOVA with Dunn’s multiple comparisons test (H-L) with significant p values indicated in the figure.

### Temporal acquisition of exhaustion phenotypes

To better understand the order in which these alterations occurred we performed a time course collecting cells after each stimulation. We quantified the progression of exhaustion by determining polyfunctional cytokine production after stimulation (IFNγ and TNFα double positive cells) (Fig. 3A). These results showed that after three stimulations the functionality of the cells decreases, and polyfunctional cytokine producing cells are virtually undetectable after the sixth stimulation. Mitochondrial mass increased and peaked initially after the second stimulation in line with the maximal cytokine polyfunctionality, an indicator of the peak effector phase. There was then a decrease in mitochondrial mass before a final increase towards the fifth stimulation as the cells entered the dysfunctional state. (Fig. 3B). ROS followed a similar trend to mitochondrial mass although the initial peak was after the third stimulation before a decrease and final terminal increase at the fifth stimulation (Fig. 3C). While high surface expression of 2B4 and CD39 was specifically observed at the later timepoints, PD1 remained highly expressed throughout the time course after activation (Fig. 3D).

**Figure 3.**
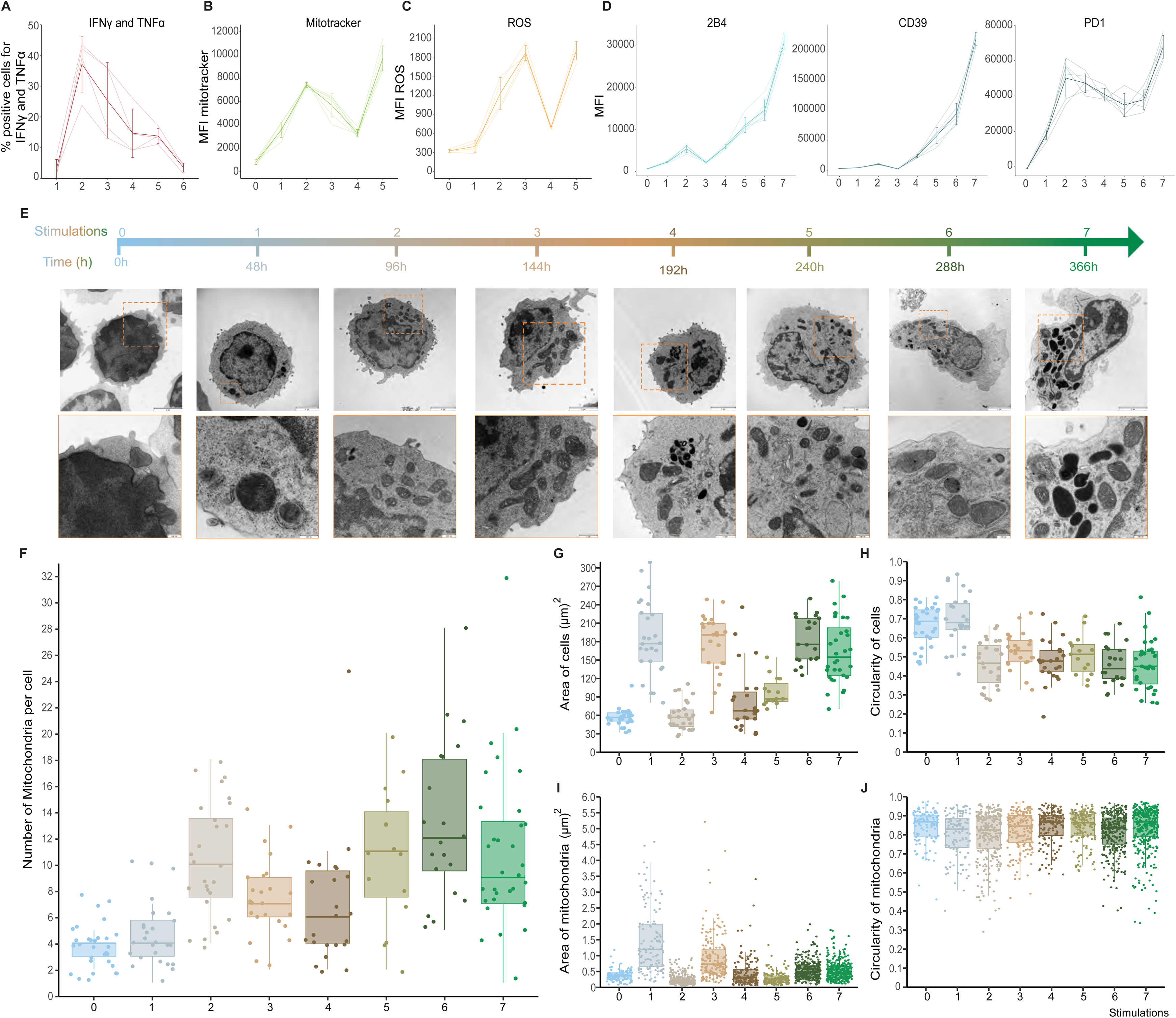
Time course of phenotypic exhaustion development. A. Percentage of cells double positive for IFNγ and TNFα after each stimulation. B. Mitochondrial mass as determined by mitotracker staining C. Cellular ROS content D. Expression of 2B4, CD39 and PD1 showing MFI after each stimulation. E. Representative TEM images of cells and inset zoom of region for mitochondria, scale as indicated on each panel. F. Number of mitochondria per cell G. Area of cells H. Circularity of cells I. Area of mitochondria J. Circularity of mitochondria Representative data from two (A-D) independent experiments, light lines individual mice and dark lines mean ± SD with n=7 mice per measurement. For TEM at least 3 sample preparations per group were imaged and 27 cells per sample were quantified. Boxplots shown at median with interquartile range with symbols representing individual cells (F,G,H) or individual mitochondria (I,J). P values for Brown-Fosythe and Welch ANOVA Test with multiple comparisons (Table S1).

We used the time course to evaluate the morphology and features of the cells and their mitochondria using TEM (Fig. 3E). We collected samples after each stimulation and evaluated size, area, number and circularity at both cellular and mitochondrial level (Fig. 3F-J). This analogous method of determining mitochondrial mass showed the same trend as was observed using mitotracker staining and quantification by flow cytometry (Fig. 3B,F, Fig 2D,E). The area of the cells had an initial cyclical expansion and contraction before maintaining an area of approximately three times that of unstimulated cells (Fig. 3G), whilst the circularity of the cells decreased from the second stimulation and remained at this reduced level for the duration of the experiment (Fig. 3H). The area of the mitochondria initially followed a similar recurring expansion and contraction aligned with the area of the cells, however, after the fifth stimulation there was no correlation and the mitochondrial area was relatively constant (Fig. 3I). There was no noticeable difference in the circularity of the mitochondria at any timepoint (Fig. 3J). These results illustrate that the phenotypic, functional and metabolic alterations observed during T cell exhaustion show marked differences in their temporal acquisition during chronic antigen stimulation.

### *In vivo* model of T cell exhaustion in response to cancer

Having established an *in vitro* model of exhaustion it was essential to compare it with *in vivo* exhausted cells. Hence, we decided to use the syngeneic model of colon adenocarcinoma, MC38, a cell line which has a moderate infiltration and responds to immunotherapy ^35^. This cell line was modified to stably express ovalbumin after genome integration using the PiggyBac transposon system. After puromycin selection this tumour cell line (MC38-OVA) was used compared to the parental (MC38) in immunocompetent mice (Fig. 4A). After the tumours formed, we identified a distinct pattern of responses to the MC38-OVA tumours. Initially, all the MC38-OVA tumours grew comparably but at a slightly slower rate to that of the parental MC38. However, after day nine it was possible to separate them in two broad response categories. One group that showed continuous growth and another group that started to decrease in size. Around day 15 three different responses could be identified: 1) *regressing*: tumours that started to decrease in size from day nine and were almost eliminated; 2) *non-responsive*: tumours that grew steadily throughout the experiment; and 3) *exhausted*: tumours that initially regressed but underwent fluctuations in tumour size (Fig. 4A). In order to classify the tumour responses in an unbiased manner the data was processed to generate the inter-measurement tumour area difference, the number of measurements that were smaller or the same as the prior measurement, the sum of the measurement changes from day 10 as well as the area under the curve (AUC) for the tumour area and change in tumour area. The tumours were classified on day 15 to 17 after tumour cell injection based on the day when samples were collected for downstream analysis from the experiment or otherwise on day 17 to be consistent. With these parameters tumours were classified as *non-responsive* if the tumour continuously grew during the measurement period. A tumour was also defined as *non-responsive* if there was only one dip, the AUC for the tumour change was greater than 0.2, the sum of the tumour size change after day 10 was greater than 0.1 with the penultimate measurement equal or greater than zero, final measurement change greater than zero and the largest in the measurement period. *Regressing* tumours were defined as those that had negative tumour change measurements during the experimental window. They also had to have a final change in tumour area that was negative or the final two measurements were less than or equal to zero. The tumours could also be classed as *regressing* if the final measurement was the smallest and the sum of the tumour size changes after day 10 was less than zero. All other mice were classified as *exhausted*. These responses were independent of age, genetic background, cage, and sex (Table. S2). The emergence of three distinct responses was unexpected given the same tumour cell line was injected. To further validate this observation, we repeated the experiment in different WT colonies and in another animal facility observing the same effect in each replicate. The graph shown in figure 4A is the result of 10 pooled experiments. These patterns of tumour responses appeared in similar proportions in each experiment, with around half (∼57%) of the animals presenting complete regression and the other half being divided between exhausted (∼29%) and non-responsive (∼14%) (Fig. 4a). A subset of mice were allowed to continue until the tumour grew and reached the experimental limit or became undetectable. For these mice they were classified again at this final experimental endpoint and out of the 85 mice originally classified as *exhausted,* 17 (20%) had rejected the tumour by the end point now classified as *regressing (exhausted-to-regressing)*. There were 200 mice classified as *regressing* with 34 (17%) becoming *exhausted* with the tumour returning at a later timepoint (*regressing-to-exhausted)*. The classification of *non-responsive* was 100% accurate (Fig. S3).

**Figure 4.**
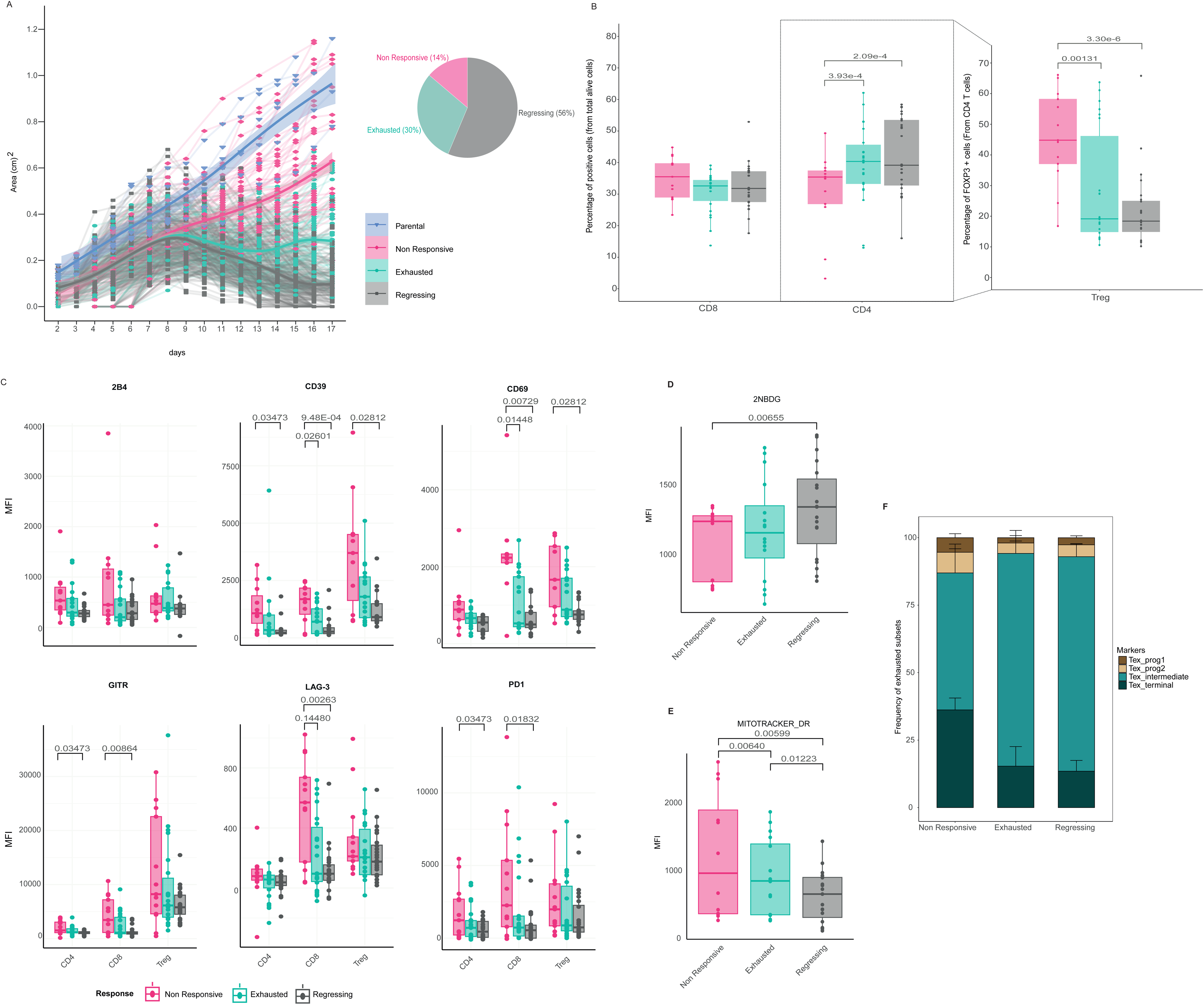
Characterization of an *in vivo* exhaustion model. A. Tumour growth of MC38 tumours in mice after injection of MC38 parental (blue) or MC38-OVA with three different responses (non-responsive in pink, exhausted in green, and regressing in grey). Average trend line in bold with 95% confidence interval with individual mice in shaded lines. B. Percentage of T cells subsets within tumours, CD4 and CD8 are expressed as percentage of alive cells and Tregs as percentage of CD4+ viable cells. C. Quantification of expression of exhaustion markers on CD4, CD8, and Tregs isolated from TILs. D. Glucose update in CD8 TILs as assessed by MFI of 2-NBDG staining. E. Mitochondrial mass of CD8s TILs from tumours determined by MFI of Mitotracker staining F. Relative proportions of exhausted subsets in CD8s-TILs Representative data from 10 pooled experiments (A), four pooled experiments (B-E), and two pooled experiments (F). Boxplots show mean and interquartile range, symbols represent individual mice. Mean ±SEM is shown (E, G) with symbols representing individual mice. Statistical analysis were performed using Phenstat with time as a fixed effect model and p values are indicated in the figures.

Considering that together with exhausted CD8 T cells, CD4 T cell subsets are the key T cell populations within the TME with implications in prognosis, we decided to look at their infiltration in the three responses. MC38-OVA cells were orthotopically implanted into FOXP3_IRES_mRFP (FIR) mice ^36^ enabling direct *ex vivo* identification without fixation and permeabilization. Immunophenotyping of tumours at 15 days post injection, showed differences in the fraction of infiltrating CD4 T cells and percentage of CD4 T cells that were Tregs. Regressing and exhausted tumours contained significantly more CD4 cells compared to non-responsive tumours, which had a higher fraction of CD4s that were Tregs (Fig. 4B). No differences were observed in draining lymph nodes or spleen (Fig. S4).

Next we determined the expression of key exhaustion related phenotypic markers that were altered in our *in vitro* model and observed similar changes in the *in vivo* model. Specifically on CD8 T cells the expression of LAG-3, CD39 and CD69 was significantly higher on the cells from non-responsive tumours compared to both exhausted and regressing tumours. There was also significantly higher levels of PD1 and GITR on CD8 from non-responsive tumours compared to regressing. A similar trend was observed for 2B4 although this was not significant. Interestingly CD39 was also significantly increased on both Tregs and FOXP3 negative CD4 in non-responsive tumours compared to regressing. PD1 and GITR were significantly increased on the FOXP3 negative CD4 in non-responsive tumours compared to regressing but not in Tregs (Fig. 4C). These changes were unique to the tumour as they were not observed in the lymph node or spleen (Fig. S4)

Next we focused exclusively on the CD8 T cells from the different responses. The glucose uptake was determined by 2NBDG staining, and the T cells from the regressing tumours had a higher uptake of glucose compared to the non-responsive (Fig. 4D). Mitochondrial mass (Mitotracker Deep Red) demonstrated a gradient effect with CD8 from non-responsive tumours the highest and regressing the lowest with exhausted in the middle significantly different from both (Fig. 4E). The frequency of exhausted CD8 subsets, defined by CD69 and Ly108, was determined and the terminally exhausted subset was higher in the non-responsive with the other responses having similar proportions (Fig. 4F). The development of this *in vivo* model with different immune responses provides a unique opportunity to study immune escape, and recurrence, and could be used to analyse other populations involved in antitumor immune response.

### Transcriptomics signatures of exhaustion *in vitro* and *in vivo*

To understand the influence of the microenvironment in the development of T cell exhaustion we performed adoptive transfer of *in vitro* activated OT-1 T cells into mice bearing exhausted tumours. Mice were subcutaneously implanted with MC38-OVA cells and a subset of mice with ‘exhausted’ tumours (started to regress but not with full clearance) were adoptively transferred with OT-1 cells after 24h of *in vitro* activation (Fig. S5A). After 21 days of *in vivo* exposure we collected tumours and draining lymph nodes, isolated the CD8 T cells, and compared the expression of characteristic exhaustion markers (PD1, 2B4, CD39 and CTLA-4) from host and transferred CD8s (Fig. S5B). There was no significant difference between the host and transferred CD8 T cell expression of the markers although for CD39, 2B4 and CTLA4 there was a trend for higher levels on the transferred CD8, perhaps due to the presence of the OVA protein within the tumour.

To fully investigate the impact of the tumour microenvironment we performed bulk RNAseq on sorted CD8 TILs from the exhausted tumours and the *in vitro* T cells (naïve, activated and exhausted). First we identified the overlapping differentially expressed genes (DEGs) that were present in the *in vitro* exhausted T cells and the *in vivo* exhausted T cells, in comparison to activated T cells, to generate the *exhausted experimental signature* consisting of 3,260 up regulated genes (Fig. 5A). Next we compared these DEGs to those identified from scRNAseq analysis of human TILs isolated from multiple cancer types ^32^ with a focus on those in the annotated CD8 exhausted clusters to generate *the exhausted signature* (Fig. 5B). This yielded 234 genes that are conserved between human and mouse and are present in both the *in vitro* and *in vivo* murine models representing key targets for therapeutic intervention. All the comparisons were performed against early activated CD8 T cells to eliminate genes that are also critical for T cell activation and results in three distinct clusters of expression patterns. (Fig. 5B). Analysis of reactome pathways via GSEA^37^ demonstrated significant enrichment of immune related pathways strengthening the assignment of these as core keys (Fig. 5B). In addition to many known exhaustion genes, including *Havcr2*, *Entpd1* and *Pdcd1,* there are many additional genes within this signature that represent novel candidates for follow up mechanistic studies.

**Figure 5.**
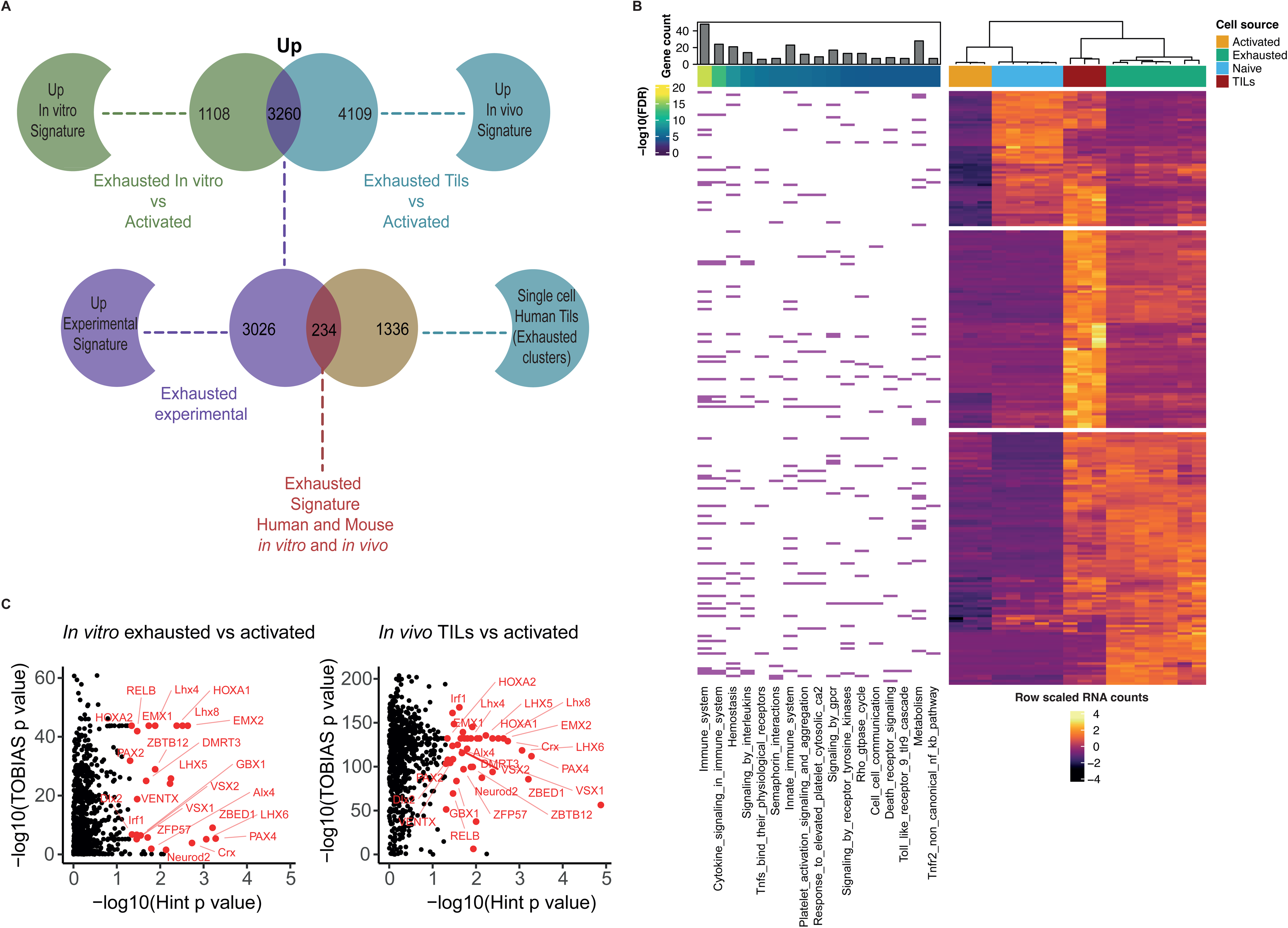
Signatures of exhaustion. A. Venn diagram of exhaustion signatures showing experimental signature and interspecific core of exhausted genes that are up regulated. B. Heat map of the conserved interspecies core of exhausted genes showing expression of naïve, activated, and exhausted *in vitro* CD8 and *in vivo* CD8 TILs for the exhausted signature. Enriched reactome pathways for genes from GSEA indicated, with FDR q-value for enrichment and number of genes per pathway. C. Enriched transcription factor motifs defined by TOBIAS and hint. Red dots indicate motifs that are significant p < 0.05 for both methods and annotations are for motifs present both in vitro (left) and in vivo (right).

Epigenetic changes have previously been reported in exhausted T cells ^6,34,38,39^. To further validate our models and to identify any potentially novel epigenetic changes in exhausted T cells, we performed an assay for transposase-accessible chromatin with sequencing (ATAC-Seq) to assess chromatin accessibility^40^. Differentially accessible peaks were annotated in the samples and transcription factor motif foot printing analysis was performed using TOBIAS and Hint ^41,42^. The transcription factor motifs that were significantly increased using both methods were compared between the *in vitro* activated and exhausted plus the *in vivo* TILs and *in vitro* activated, yielding a total of 24 that were shared in both the *in vitro* and *in vivo* exhausted CD8 T cells (Fig. 5C). Interestingly, HOXA2 was previously shown to be enriched in exhausted T cells from the LCMV model^43^. Many of other transcription factors represent novel areas for further study.

## Discussion

A greater knowledge of the dynamics of T cell exhaustion is crucial to enhance the efficiency of future immunotherapies. However, developing a deeper understanding of the molecular, morphological, and functional characteristics of exhausted T cells is hampered by the lack of a reliable method to produce these cells for study in tractable *ex vivo* models. Here we present an *in vitro* model that allows direct comparison with TILs from both *in vivo* murine models and human studies. This model is a fast and reproducible system to study exhaustion in the laboratory by chronic antigenic stimulation in the absence of any other components of the tumour microenvironment. Other models of T cell exhaustion specific to viral infection ^44^ and non-specific TCR stimulation mediated by antibodies^45^ have emerged as a useful tool to perform CRISPR based screenings, as alternatives to *in vivo* models. However, they have not characterised the temporal acquisition or diversity of the features of exhaustion that we have undertaken. The model presented here demonstrates the reproduction of the hallmarks of exhaustion; high expression of IRs, decreased proliferation, decreased cytokine production and the alteration of metabolic and mitochondrial function ^1,6,19,20^, confirming that this is a reliable system to study exhaustion. Temporal analysis allowed the identification of the different phases in the development of exhaustion, and alluded to the sequential order of the alterations present in this population. The TCR strength of the OT-1 used in this model plays a fundamental role as it produces a strong signalling cascade associated with a faster induction of exhaustion ^46^, while still being a suitable model to study the specific response to a cognate antigen in the context of cancer.

It is established that upon T cell activation, metabolic alterations result in increased mitochondrial mass and ROS ^47,48^ and exhausted cells exhibited a further increase in both when compared with activated cells. Exhausted cells had a higher uptake of glucose and a decrease in the production of IFNγ, TNFα and CD107a, highlighting a dysfunctional state. The importance of mitochondrial activity in exhausted cells has been shown previously ^49,50^. Here we demonstrate the sequential nature of these events within the exhausted cells, with changes in the morphology of mitochondria and cells. These changes are linked to the functional and metabolic state of the cells. A similar relationship between metabolic stress and altered mitochondrial morphology has previously been demonstrated with hypoxic cells ^8,9^. We noticed that after the third stimulation it was possible to observe electron-dense structures that looked similar to a mitochondrial-lysosome-related organelle (MLRO) that had been described in hepatocytes dedifferentiation^51^. However, more characterization is needed in these structures for exhausted T cells.

Our *in vivo* model allows the evaluation of different immune responses against the same tumour model expressing the OVA antigen, and permits the exploration of the immune populations involved in different responses. As in our model, a high infiltration of regulatory T cells (CD4+ FOXP3+) and increased CD4s have been shown to be present in non-responsive and regressing human tumours respectively ^52–55^, demonstrating the validity of our model. The comparison of *in vitro* and *in vivo* exhausted cells allows the identification of drivers of the exhaustion phenotype that are shared *in vitro* and *in vivo* and which are the genes that are specific to each condition. TILs from non-responsive and exhausted tumours expressed higher levels of key markers of exhaustion and increased mitochondrial mass when compared to the TILs from the regressing tumour response. These differences in mitochondria and exhaustion markers *in vivo* were similar to our observations *in vitro* when comparing exhausted and activated T cells. The high potential of the mitochondria has been associated with a dysfunctional state previously both *in vitro* and *in vivo* ^56,57^. However, other studies have associated the dysfunctional state with depolarized mitochondria ^50^ in exhausted cells. This discrepancy could potentially be explained by differences in experimental techniques, our measurements were direct using TMRE while other studies have utilized mitotracker stains. TILs from the regressing tumours had increased glucose uptake compared with TILs from the non-responders and exhausted tumours. In contrast, the exhausted cells *in vitro* had a higher glucose uptake compared with activated cells. This differential response could be due to the differences between the metabolic requirements of the TILs and the availability of glucose within tumour microenvironment.

As many of the differentially regulated genes in exhaustion can also be present in activated T cells all comparisons were made against freshly activated T cells, this allows the identification of exhaustion specific drivers rather than those that play a role in T cell activation. One of the possible limitations of the *in vitro* model is the lack of any other components of the TME and the only driver for the induction of exhaustion being chronic TCR signalling. However, given the overlap from the RNAseq of 3,260 upregulated genes there is a core set that are common between the *in vitro* and *in vivo* models and represent factors that are independent of the TME. Genes specifically affected by the TME would be present in the 4,109 up regulated genes that are unique to the *in vivo* model with a further 1,108 genes only found *in vitro*. To further support the validity and translational potential of the model we identified that 234 genes present in our core experimental signature are also enriched within exhausted CD8 T cells isolated from a panel of human tumours^32^. The genes found in the exhausted signature contained well established genes previously linked to exhaustion, including: *Pdcd1*, *Havcr2* and *Entpd1*. As well as those identified more recently through CRISPR screenings including *Bhlhe40* ^44^, and single cell CRISPR screenings *in vivo* including *Rbpj* ^58^.The latter two examples are transcription factors that promote differentiation of exhausted T cells. Epigenetic changes are also critical in the development and maintenance of the exhausted state ^6,34,38,39^ and we have identified 24 transcription factor motifs that have increased accessibility in both the in vitro model and in vivo TILs. This includes HOXA2, which has previously been linked to virally exhausted T cells ^43^, with many others representing new avenues to investigate their role in T cell exhaustion. In summary we present an *in vitro* model and key datasets that could be utilised in the development of targets for T cell reinvigoration in cancer immunotherapy.

## Material and methods

### Mice

Wildtype mice were obtained from genetically altered C57BL/6NTac lines from the Wellcome Sanger Institute Mouse Genetics Project or Taconic. OT-1 (C57BL/6-Tg(TcraTcrb)1100Mjb/J, RRID:IMSR_JAX:003831) and FIR (C57BL/6-*Foxp3^tm1Flv^/*J, RRID:IMSR_JAX:008374) mice were obtained from The Jackson Laboratories. Animals were housed in Sanger Institute Research Support Facility and University of Cambridge Biological Services Unit under Specific Opportunist Pathogen Free conditions in Individually Ventilated Cages at a density of 1 to 6 animals per cage. Mice had ad libitum access to irradiated or autoclaved diet and autoclaved water with aspen bedding and environmental enrichment provided. The care and use of all mice in this study were in accordance with the Home Office guidelines of the UK and procedures were performed under a UK Home Office Project License (P2E57E159) which was reviewed and approved by the Sanger Institute’s and University of Cambridge Animal Welfare and Ethical Review Body. For *in vitro* and *in vivo* experiments mice were between 7 and 20 weeks old at experimental initiation. This was a multigenerational study with cohorts of sizes between 20 and 70 animals size dependent on sex and age matching and expected dosing groups to achieve sufficient replicates of each result. Where possible, age and sex matched mice in the cohorts were littermates and both genders were used to reduce sex bias.

### Reagents

Unless otherwise stated tissue culture reagents and other biochemicals were from Sigma-Aldrich and ThermoFisher.

### *In vitro* exhaustion protocol

CD8 T cells were isolated from spleens of OT-1 mice using the CD8 mouse isolation kit (Miltenyi Biotec) according to the instructions of the manufacturer. Cells were counted and seeded at a density of 1×10^6^ per mL in complete media: RPMI 1640 supplemented with 10% heat inactivated fetal bovine serum (FBS), 50 μM 2 mercaptoethanol, 1% non-essential amino acids, 1 mM sodium pyruvate, 10 mM HEPES and 1% L glutamine, penicillin, streptomycin (CM). The OVA peptide SIINFEKL (AnaSpec) was added at 3 μg/mL with 100 U/mL human IL-2 (Peprotech). Every 48 hours the cells were centrifuged and the pellet was resuspended between 1 and 2 × 10^6^ per mL in fresh complete media containing the OVA peptide 3 μg/mL and IL-2 (100 U/mL) after the second stimulation IL-15 (10 ng/mL, Peprotech) was added The number of stimulations is indicated in the figure/text. Comparisons were made with CD8 T cells activated for 24h with OVA (3 μg/mL) in CM with 100U of IL-2 and expanded for 72h if indicated.

### Proliferation with 5-Ethynyl-2-deoxyuridine (Edu)

Cells were incubated with EdU (10 μM) for one hour in CM, then stained for viability with the live/dead fixable red stain according to manufactures instructions. Cells were fixed with 4% PFA for 15 minutes at room temperature. Then washed with permeabilization buffer from the Foxp3/Transcription Factor Staining Buffer Set (eBioscience™) and incubated in 10 mM sodium ascorbate, 2 mM copper sulfate and 1 μM Alexa647 azide dye for 30 minutes at room temperature. After washing, cells were labelled with 2 μg/ml Hoechst 33452 for 15 minutes. Cells were then analysed by flow cytometry on a Fortessa SORP instrument (BD Biosciences).

### Preparation of tissue samples for sorting and FACS

Tumours were collected and processed using the tumour dissociation kit (Miltenyi Biotech) and GentleMACS C tubes using a GentleMACS dissociator according to the supplied protocol for soft//medium tumours. T cells from tumour single cell suspensions were enriched with the CD8/CD4 (TIL) MicroBeads kit according to the manufacturer’s instructions (Miltenyi Biotech). Lymph nodes and spleens were crushed through a 70 μm filter (MACS Smart Strainer) and washed with PBS. Cells from spleens were treated with red blood cell lysis, centrifuged and washed with PBS.

### Flow sorting

After tissue processing to single cell suspension, cells were stained with CD8a and eBioscience™ Fixable Viability Dye eFluor™ 780. Cell sorting was performed on a BD FACS Aria with a 100 μm nozzle.

### Flow cytometry

For surface membrane markers cells were resuspended in titrated amounts of antibodies (Table S3) diluted in FACS buffer (D-PBS without calcium and magnesium, 0.5 % FBS, 2 mM EDTA and 0.09 % sodium azide) and incubated in the fridge for 30 minutes. Viability was determined using a compatible dye (eBioscience™ Fixable Viability dye eFluor™ 780, eFluor™ 506, eFluor™ 455UV or DAPI). Where required for intracellular staining cells were fixed with an appropriate fixative either Foxp3/Transcription Factor Staining Buffer Set (eBioscience™) or Cytofix/Cytoperm (BD) and intracellular staining performed in the appropriate permeabilisation buffer (either for 45 min or overnight in the fridge). Samples were analysed on a MACSQuant VYB, BD Fortessa or Cytex Aurora. Compensation samples were prepared with Ultracomp eBeads (eBioscience™), Arc reactive beads (Invitrogen™), anti-REA Comp beads (Miltenyi Biotec) and unstained cells and determined on the instrument using inbuilt software. Data was analysed with FlowJo software (v10.7.1, BD) using singlets and viable cells for the in vitro experiments and for the in vivo experiments using the gating strategy outlined in supplementary figure 6.

### Degranulation assays to evaluate cytokine production

Cells were seeded in a 96 well plate with CM media containing the protein transport inhibitor monensin (BD GolgiStop™, 1 in 1000) and CD107a antibody (0.75 μg/ml), with or without OVA (2 μg/mL) for four hours at 37°C in 5% CO_2_. Cells were stained with eBioscience™ Fixable Viability dye eFluor™ 780 and fixed with Cytofix/Cytoperm solution for 20 minutes at 4°C. After washing with perm/wash solution (BD, USA) cells were stained with antibodies to cytokines (Table S2) and samples acquired on a flow cytometer (MACSQuant VYB, BD Fortessa or Cytek Aurora). Data was analysed in FlowJo.

### Metabolic stress test

Exhausted and activated T cells were seeded on a poly-D-lysine-coated 96-well plate (200,000 cells per well). T cell fitness standard kit test (Agilent, Santa Clara, CA) was performed on a Seahorse XFe96 Analyzer. Metabolic rate was normalized to cell count determined on a Moxi (Orflo).

### Mitochondria and metabolism profiling through FACS

Cells were seeded in 96 well plates and stained using: reactive oxygen species (CellROX™ Deep Red Reagent, 5 μM), mitotracker (MitoTracker™ Deep Red FM, 2 μM) and TMRE (Tetramethylrhodamine ethyl ester, 1 μM) in complete media. To measure glucose uptake cells were seeded in growth media without glucose with the addition of 20 μM 2-NBDG (2-(*N*-(7-Nitrobenz-2-oxa-1,3-diazol-4-yl)Amino)-2-Deoxyglucose). Cells were incubated for 30 minutes at 37°C in 5% CO_2_ then washed and stained with DAPI (1:5000) for viability and acquired on a flow cytometer (MACSQuant VYB, BD Fortessa or Cytek Aurora). Data was analysed in FlowJo.

### Confocal microscopy

Cells were seeded in bottom glass dishes and stained with nucblue (2 drops per 1 mL of NucBlue™ Live ReadyProbes™ Reagent (Hoechst 33342) ThermoFisher) and mitotracker (MitoTracker™ Deep Red FM, 2 μM ThermoFisher). Cells were acquired in a confocal microscope Leica SP8 and processed with the Leica Application Suite.

### Electron microscopy

T cells were pelleted and fixed (using a solution of 2.5% glutaraldehyde and 2% paraformaldehyde in 0.1 M sodium cacodylate buffer) at isolation (Naïve) and every 48 hours after stimulation. Cells were stained with 1% osmium tetroxide and mordanted with 1% tannic acid, followed by dehydration using an ethanol series (contrasting with uranyl acetate at the 30% stage) and embedding with an Epoxy Resin Kit (Sigma-Aldrich). Ultrathin sections were cut on a Leica UC6 ultramicrotome contrasted with lead citrate and images were acquired on a FEI Spirit Biotwin, using a Teitz FX416CCD Tem Cam. Analysis of EM images to determine, area and circularity of cells and mitochondria was done using the open source project for image analysis ImageJ, also known as Fiji following the methods described by Lam et al.^59^ Fourteen random samples were blinded measured and counted by three independent researchers to confirm results. The formula for circularity is 4π(area/perimeter^2). A value of 1.0 indicates a perfect circle.

### *In vivo* experiments

MC-38 OVA cells were generated by inserting the cOVA sequence from PCI-neo-cOVA (a gift from M. Castro, RRID:Addgene_25097), into PB-CMV-MCS-EF1a-Puro PiggyBac vector (PB501b-1, System Biosciences) to generate PB-CMV-cOVA-EF1a-Puro plasmid which was delivered together with piggybac transposon (a gift from A. Bradley) to MC-38 cells (a gift from L. Borsig). After selection with puromycin (2 μg/ml, Invivogen) expression of OVA was determined by western blotting (Abcam) and presentation of SIINFEKL peptide in the context of MHC Class I on the surface of MC-38 cells was determined by flow cytometry (H-2K^b^/SIINFEKL, REA1002, Miltenyi Biotec). The MC-38 cell line was maintained as a pool. Wild type or FIR mice received between 1 and 2.5×10^6^ MC-38 OVA or MC-38 parental cells subcutaneously (exact cell numbers for each cohort are detailed in the supplementary metadata Table S2).

Tumour measurements were collected at regular intervals by one individual using a caliper and tumour area calculated (length x width (cm)). Tissue collection was performed on days 15, 16 and 17 (Table S2) and the cohort of mice was classified on the day of tissue collection or day 17 whichever was sooner. From the tumour areas the following parameters were derived in R, maximum area day, minimum area day, inter-measurement difference, number of measurements the area remained the same, number of measurements the area decreased compared to the prior measurement, sum of the inter-measurement difference after day 10, and area under the curve of changes in tumour area after day 10. In addition the tumour area change on the final day and penultimate measurement were used to classify the tumours as outlined in the text and supplementary script. For adoptive transfer experiments WT mice were injected subcutaneously with MC-38 OVA 2 ×10^6^ cells. On day 16 post tumour cell administration OT-1 cells that had been activated *in vitro* with OVA (2 μg/mL) for 24 hours prior were injected via tail vein injection (5 × 10^6^ cells/mouse) and tissues harvested 21 days after T cell transfer.

### RNAseq

Cells were processed in Trizol (Life Technologies) and total RNA was isolated using the RNA Clean and Concentrator (Zymo Research). Libraries were prepared using the Illumina stranded total RNA prep with Ribo-Zero plus kit according to the manufacturer’s instructions and sequenced on an Illumina HiSeq 4000 with 75 bp paired end reads aiming for > 30 million reads per sample. Gzipped FastQ files were aligned to GCRm38 using star 2.7.9a ^60^, called via rsem 1.3.1^61^ (paired end mode, all other settings as default) to quantify transcript level counts. Differential expression was assessed using the R package DESeq2^62^ in R-4.1.0^63^ genes assigned a FDR<= 0.01 and displaying a log fold change of at least 2 were said to be called significant. Reactome pathway enrichment was performed on the GSEA ^37^ website (MSigDB 2024.1) using a FDR q value of 0.0001 as the threshold.

### ATACseq

10,000 sorted cells were treated as previously described^40^ using the Illumina Tagment DNA TDE1 Enzyme and Buffer Kits according to the manufacturer’s instructions. DNA was cleaned using the Monarch kit (NEB) and quantified using a bioanalyzer (Agilent). Nextera adapters were added using Nextera XT DNA library preparation kit with 9 PCR cycles performed. Samples were sequenced on an Illumina HiSeq 4000 with 50 bp paired end reads. FastQ files were aligned to GRCm38 using bwa 0.7.17 using the following flags mem -t 16 -p -Y -K 100000000. A PhiX alignment was also performed for filtering purposes and duplicate reads were marked using biobambam2 (2.0.79) and samtools. Peaks were called by analysing mapped bam files with macs2 callpeak MACS 2.2.6.1 ^64^, with bigwig support enabled (i.e. with the - - bdg flag), a q value of 0.05 and –scale-to set to small. Footprint calling was performed using HINT (1.0.2) from the rgt package, more specifically “rgt-hint footprinting”, followed by “rgt-motifanalysis matching” and finally “rgt-hint differential”. Footprint analysis was performed using TOBIAS 0.17.0 via “TOBIAS ATACorrect”, followed by “TOBIAS FootprintScores” and finally “TOBIAS BINDetect”.

### Statistical analysis

Statistical comparisons were performed using Prism 10.0 (GraphPad) with the tests outlined in the figure legends. *In vivo* experiments data were analyzed with Phenstat in R using the time as a fixed effects model to allow pooling across multiple experiments with the package Kurbatova N, Karp N, Mason J, Haselimashhadi H (2023). *PhenStat: Statistical analysis of phenotypic data*. doi:10.18129/B9.bioc.PhenStat 2.38.0, https://bioconductor.org/packages/PhenStat.

## Acknowledgments

We thank Dr Simon Clare and Katherine Harcourt for technical support. We acknowledge the services provided by the NIHR Cambridge BRC Cell Phenotyping Hub, as well as the Cytometry Core Facility, DNA pipelines and Informatics at the Wellcome Sanger institute (supported by Wellcome Grant 206194). We are grateful for the facilities and support provided by the Wellcome Sanger Institute Research Support facility and University of Cambridge University Biological Services Unit. We thank Dr Holly Robertson, Dr Victoria Harle, and Dr Vincent Zecchini for their careful reading and helpful comments on the manuscript and Dr Juan Pablo Narvaéz Goméz for his help with code and independent measurements of mitochondria. AMR was supported by a grant from Open Targets (grant OTAR2049), BF and AOS were supported by a package from Wellcome Sanger Institute and Wellcome Trust. DAG was supported by the Wellcome Trust, SH was supported by Open Targets and DJA was supported by the Wellcome Trust.

## Author contributions

AOS funding acquisition, help with experiments and analysis, reviewing and editing text. AMR: performed all the experiments, data analysis, writing and editing paper. BF: help with experiments and analysis. SH bioinformatics analysis of RNA and ATAC sequencing. DAG generation of EM and confocal imaging data. DJA funding acquisition, editing of text

## Data availability

The RNA sequencing data is available from the European Nucleotide Archive under accession: ERP161925 and the ATAC sequencing under accession: ERP124831. All other data is available in the Supplementary information.

## Supplementary information

### Tables

**Supplementary table 1:** Results of the ANOVA tests with multiple comparisons for the Figure 3; F-Mitochondria count, G-Area of cells, H-Circularity of cells, I-area of mitochondria, J-Circularity of mitochondria.

**Supplementary table 2:** Metadata and tumour measurements of all *in vivo* experiments (two sheets according to tumour cell line).

**Supplementary table 3:** Antibodies for flow cytometry used

### Figures

**Figure S1. Exhausted subsets found *in vitro*.**

Representative exhausted subsets as defined by CD69 and Ly108 expression in naïve, activated and exhausted T cells samples *in* vitro. Mean SE, n=6.

**Figure S2. ECAR profile**

Representative ECAR trace of exhausted and activated cells, Mean and SEM error, n=4

**Figure S3. Tumour growth**

A. Tumour growth curves of all animals up to final day showing the mean of their response, including mice that had a changed on their initial response from exhausted to regressing and from regressing to exhausted.

B. Tumour growth curves separated by response

Average trend line in bold with 95% confidence interval with individual mice in shaded lines. Red dashed lines represent the classification window.

**Figure S4. Lymph node and spleen phenotyping of immune responses**

A. Total CD4 or CD8 T cells (as % of alive cells) or Tregs (as % of alive CD4 cells) in spleens and lymph nodes of tumour bearing mice separated by response. Symbols represent individual mice.

B. MFI of markers associated with exhaustion on CD4 (Foxp3-), CD8 and Tregs (CD4 Foxp3+) in spleens and lymph nodes of tumour bearing mice separated by response. Symbols represent individual mice.

**Figure S5. *Tumour growth* in cell transfer experiment**

A. Tumour growth showing all mice from the experiment and in dark green tumours used for sorting of CD8 cells.

B. Expression of 2B4, CD39 and CTLA-4 of host and transferred CD8s TILs.

**Figure S6. Flow cytometry *g*ating strategy for tissue samples**

A. Identification of CD8 and CD4 total, plus CD4 subsets (CD4 Foxp3- and CD4 Foxp3+) from mice.

B. Identification of CD8 and OT-I T cells from adoptively transferred mice.

