## Supplementary figures and images for "Simulating CD8 T Cell Exhaustion: A Comprehensive Approach"

### Supplementary figures

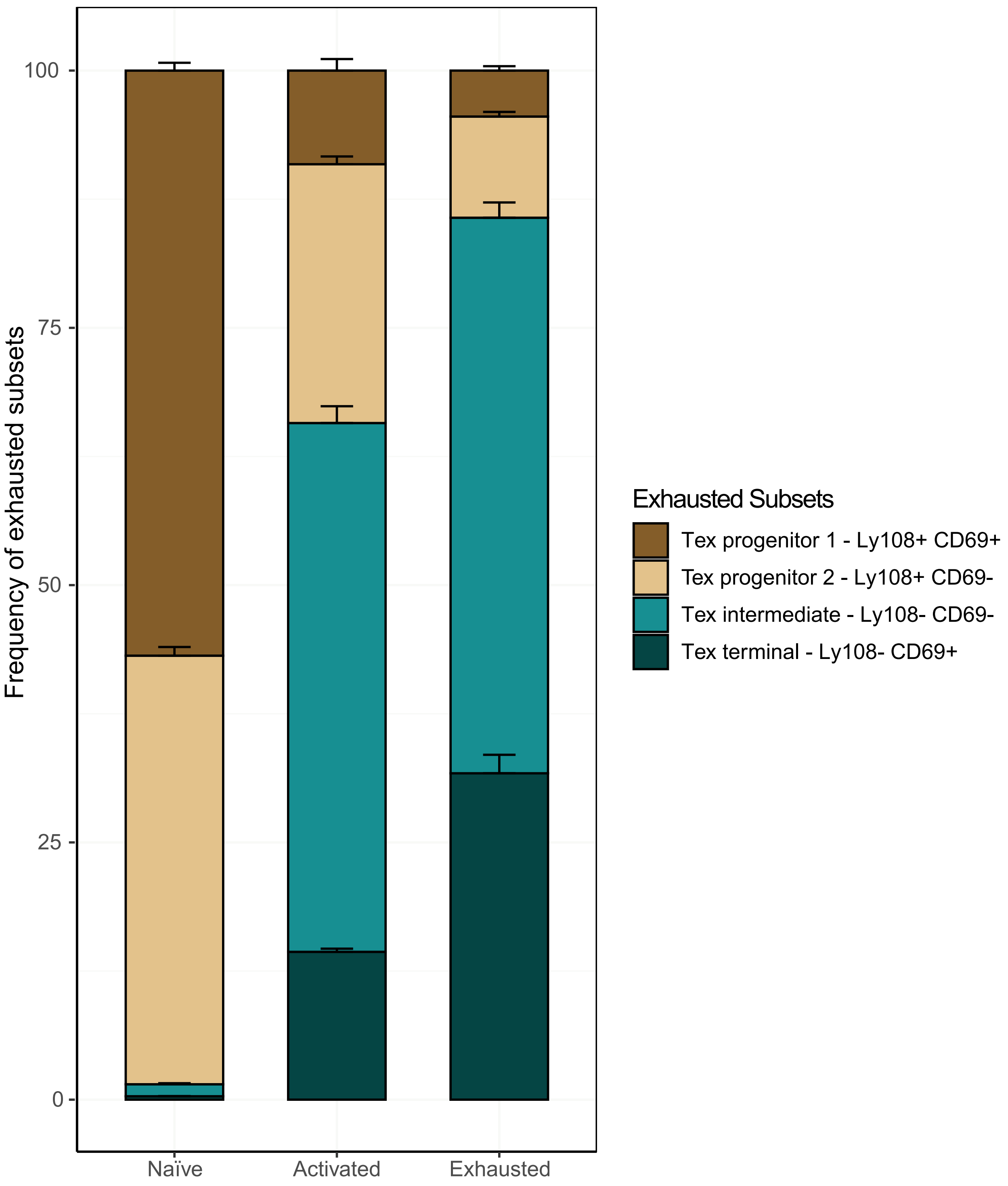

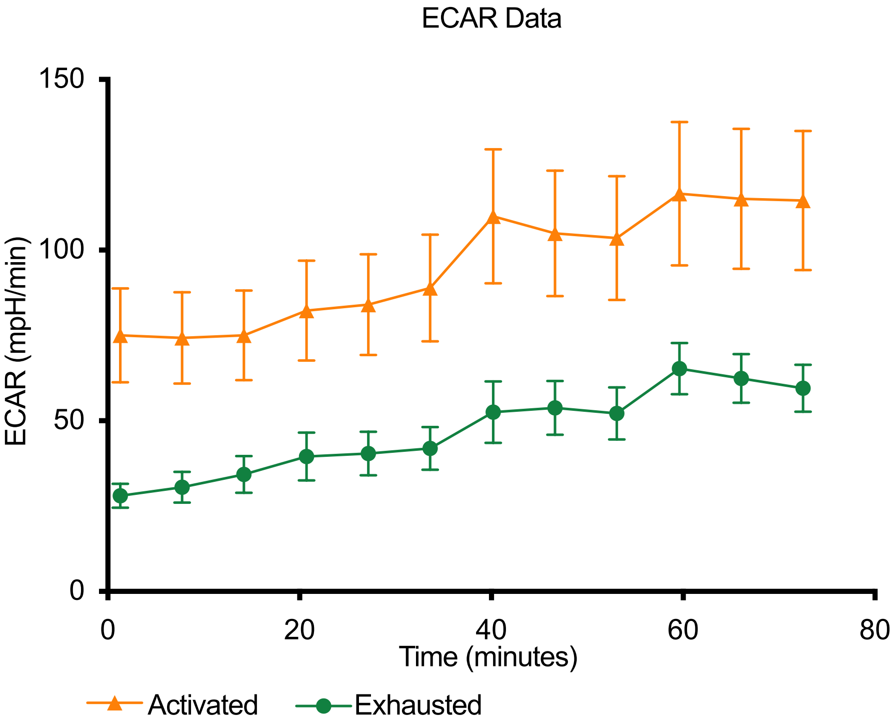

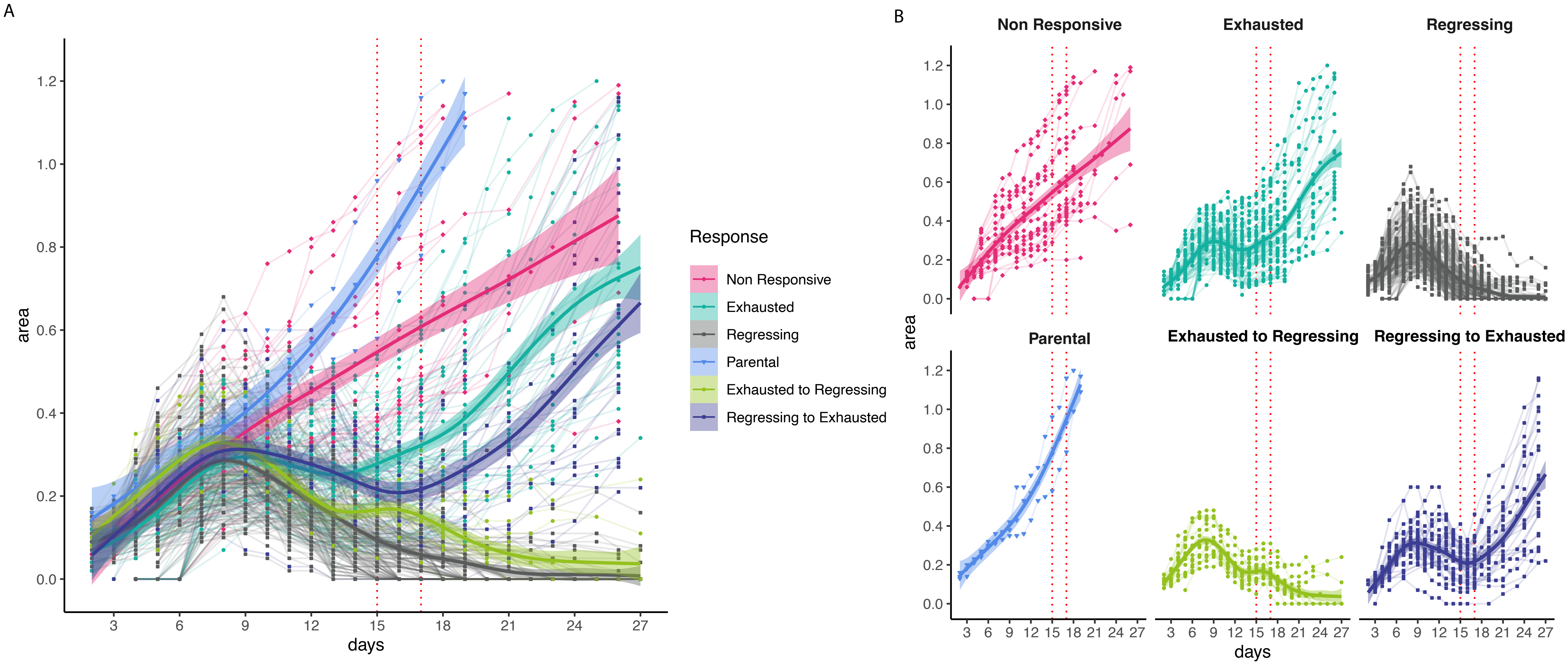

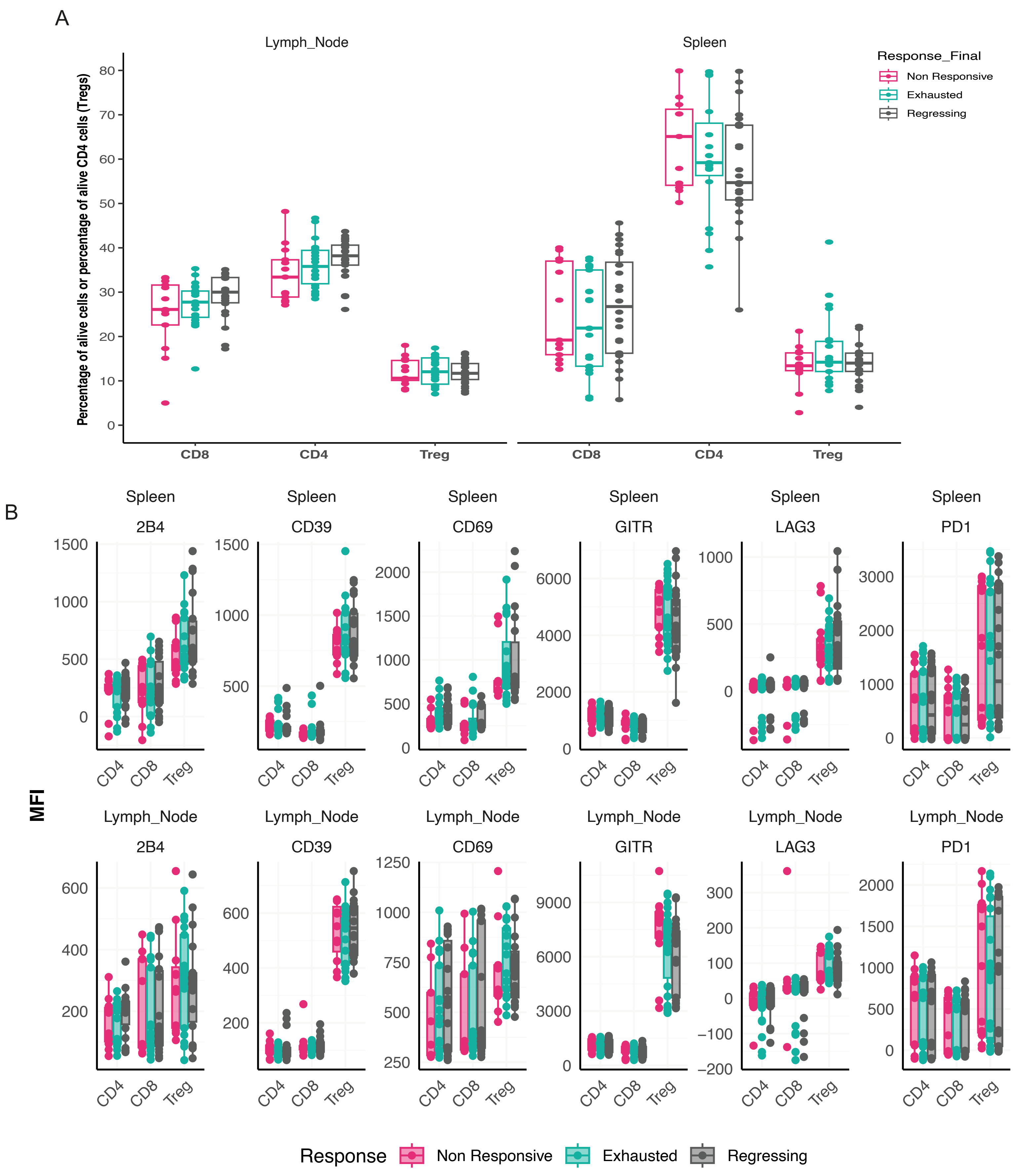

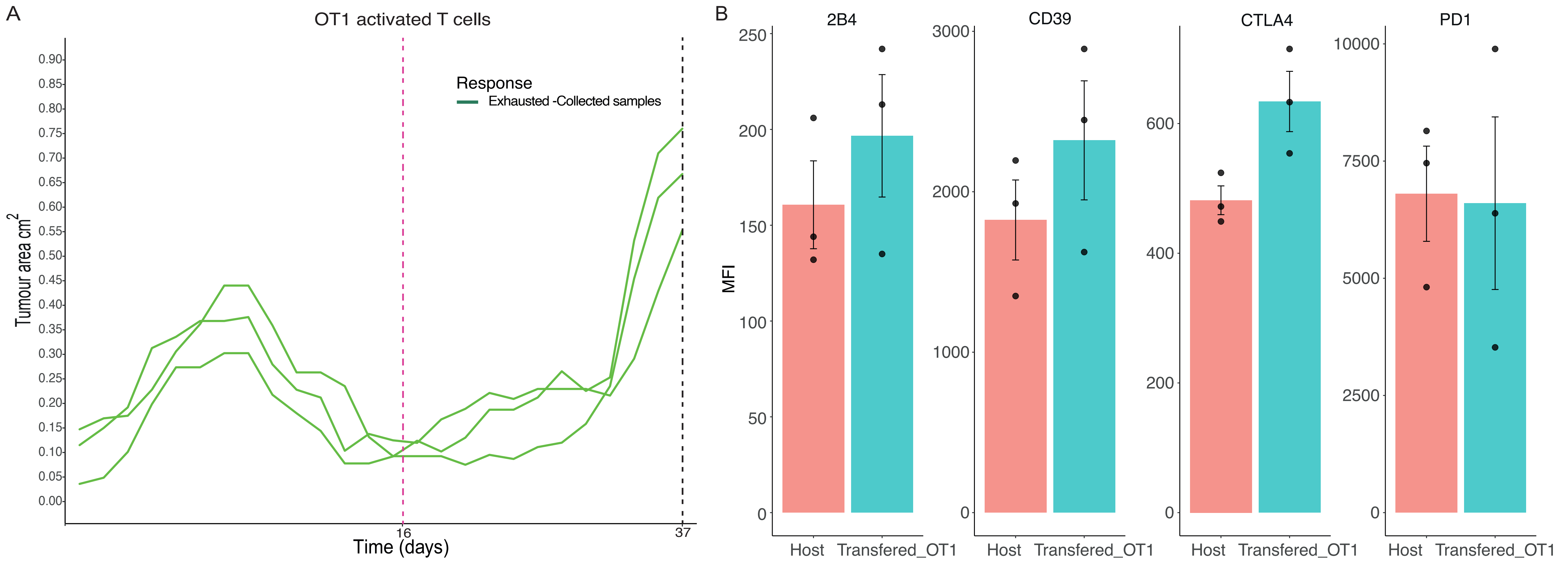

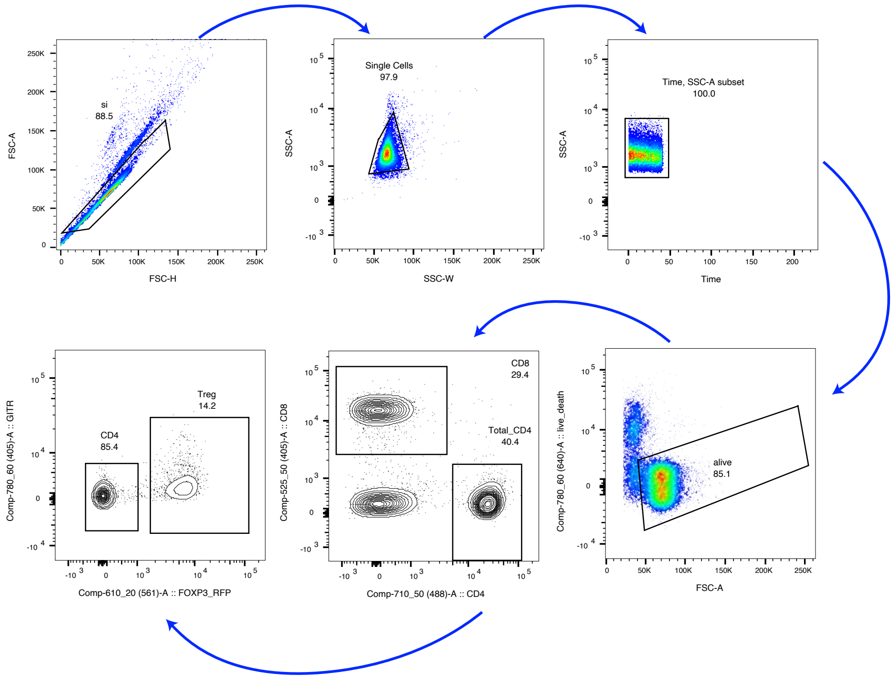

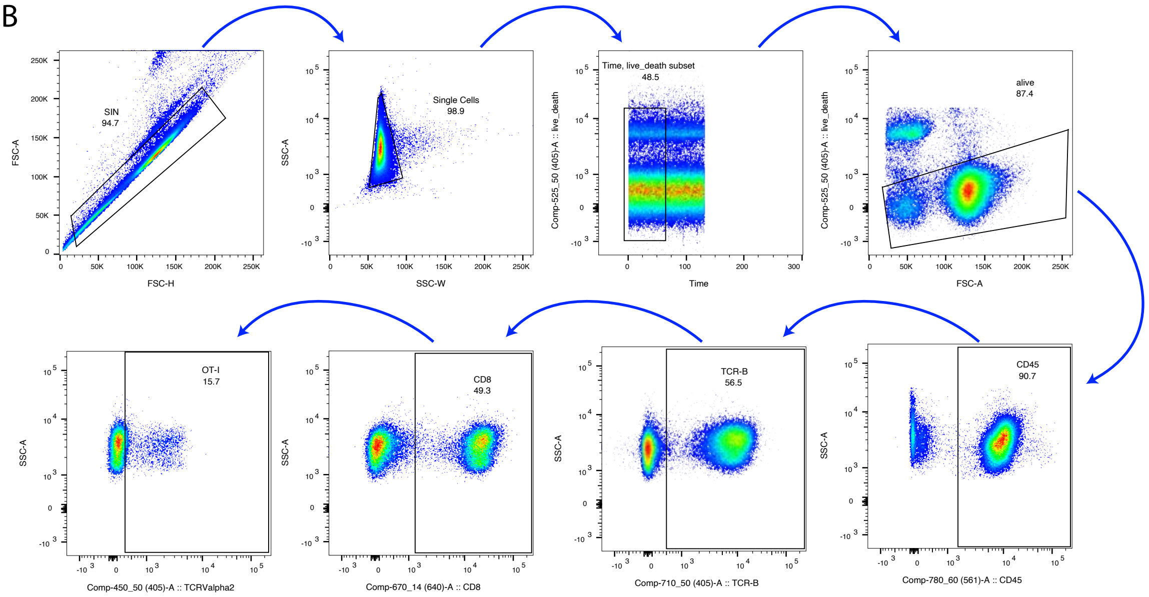
